# Rice annexin OsANN4 interacting with OsCDPK24, reduces root aerenchyma formation by modulating H_2_O_2_ under ABA treatment

**DOI:** 10.1101/519199

**Authors:** Qian Zhang, Tao Song, Can Guan, Yingjie Gao, Jianchao Ma, Zhiguang Qi, Jingze Liu, Ying Zhu, Zhengge Zhu

**Affiliations:** Hebei Key Laboratory of Molecular and Cellular Biology, Key Laboratory of Molecular and Cellular Biology of the Ministry of Education, College of Life Science, Hebei Normal University, Hebei Collaboration Innovation Center for Cell Signaling, Shijiazhuang, Hebei, 050024, China; The Institute of Viral and Biotechnology, Zhejiang Academy of Agricultural Sciences, Hangzhou, 310021, China

**Keywords:** Annexin, Calcium-dependent protein kinase, Phosphorylation, H_2_O_2_, Abscisic acid, *Oryza sativa*

## Abstract

Plant annexins are calcium- and lipid-binding proteins that have multiple functions, and a significant amount of research on plant annexins has been reported in recent years. However, the functions of annexins in diverse biological processes in rice are largely unclear. Herein, we report that *OsANN4*, a calcium-binding rice annexin protein, is a substrate for OsCDPK24, and the OsANN4 phosphorylation site is the 13th serine, which is a key site for phosphorylation. Most strikingly, abscisic acid (ABA) promotes the interaction between OsANN4 and OsCDPK24. Moreover, knocking down *OsANN4* by RNA interference resulted in visible and invisible phenotypes with exogenous ABA treatment, such as shorter shoots of seedlings, less lateral roots, earlier root aerenchyma formation and so on. The further analyzed results showed that decreased superoxide dismutase (SOD) and catalase (CAT) activity of the RNAi lines, which control H_2_O_2_ accumulation for redox homeostasis, and further promoted earlier aerenchyma formation of the root. These results suggest that a proposed molecular mechanism exists between OsANN4 and H_2_O_2_ production to response ABA.

**Highlight:** *OsANN4* enhances SOD and CAT activities to scavenge H_2_O_2_ to alleviate the formation of aerenchyma under ABA treatment. OsCDPK24 interacts with and phosphorylates OsANN4, this interaction strengthened under ABA treatment.

## Introduction

Abscisic acid (ABA), a well-known long-distance signaling molecule utilized for communication between plant roots and shoots under water-deficient conditions, is also considered a hormone that plays a critical role in abiotic stress tolerance in plants (Pekic *et al*., 1995; Cutler *et al*., 2010). Recently, many mediators involved in ABA signaling, such as ABA receptors (McCourt and Creelman, 2008; Klingler *et al*., 2010; He *et al*., 2014) and targets of ABA receptors (Bueso *et al*., 2014; Ye *et al*., 2017), have been characterized. Since the identification of the steroidogenic regulatory protein (StAR)-related lipid-transfer (START) domain as a candidate ABA receptor (Ma *et al*., 2009; Santiago *et al*., 2009; Nishimura *et al*., 2010), pyrabactin resistance 1 (PYR1) and PYR1-like 1-13 (PYL1-PYL13) have been considered as key components of the core ABA signaling pathway (Fujii *et al*., 2009; Ma *et al*., 2009; Park *et al*., 2009; Zhao *et al*., 2013). ABA receptors may inhibit the activity of phosphatase 2Cs (PP2Cs) and result in the activation of Suc nonfermenting-1-related protein kinase 2 (SnRK2s), further promoting the expression of some downstream transcription factors or membrane ion channels (Melcher *et al*., 2010; Ng *et al*., 2014).

Exposure to abiotic stresses, such as salt, drought and oxidative stress, adversely affects plant growth and crop productivity. ABA functions as an important phytohormone that has long been known to play a critical role in stress responses (Giraudat *et al*., 1994; Himmelbach *et al*., 2003) to regulate the expression of many genes, leading to complex physiological and metabolic responses that enable plants to confer tolerance to abiotic stress (Cutler *et al*., 2010; Umezawa *et al*., 2010). Increasing evidence shows that ABA-enhanced abiotic stress tolerance might be associated with the induction of antioxidant defense systems, including reactive oxygen species (ROS) scavenging enzymes, such as catalase (CAT), superoxide dismutase (SOD), ascorbate peroxidase (APX) and non-enzymatic antioxidants (Jiang and Zhang, 2002; Miao *et al*., 2006; Neill *et al*., 2008; Xing *et al*., 2008; Miller *et al*., 2010; Ozfidan *et al*., 2012; Ding *et al*., 2013). Especially the recent study shows in the presence of ABA, the increased production of H_2_O_2_ activates a calcium-dependent protein kinase (Ni *et al*., 2018), it suggests ABA induced H_2_O_2_ production to play a key role in response to abiotic stress.

Annexins belong to an evolutionarily conserved multi-gene protein superfamily comprising Ca^2+^-dependent phospholipid-binding proteins. Calcium ions bind to annexin molecules mainly via coordination sites termed type II (“AB”) high-affinity Ca^2+^-binding sites. In addition, annexins may contain two other binding sites with lower affinity, named the type III “B” and “DE” sites (Moss and Morgan, 2004). Some annexins have no type II Ca^2+^-binding sites, including some rice annexins, indicating that different protein conformations and different mechanisms for binding phospholipids may exist.

Plant annexins are reportedly tissue-specific, and their expression is regulated developmentally. Recent results also suggested that annexins play an important role in plant stress responses (Clark *et al*., 1998; Lee *et al*., 2004; Clark *et al*., 2012; Jami *et al*., 2012; Richards *et al*., 2014; Qiao *et al*., 2015), and several plant annexins have been demonstrated to play specific roles in ABA treatment and osmotic stress. The alfalfa (*Medidago sativa*) annexin gene *AnnMs2* was first reported to be activated by drought (Kovacs *et al*., 1998). *AtANN1* and *AtANN4* play important roles in osmotic stress and ABA signaling in a Ca^2+^-dependent manner (Jami *et al*., 2012). *AtANN1* was proven to act as an H_2_O_2_ sensor (Laohavisit *et al*., 2010). The expression of annexin upon ROS regulation have been identified in several organism types, including plants and animals (Richards *et al*., 2014). The rice *OsANN1* gene shows the ability to confer heat and drought stress tolerance by modulating antioxidant accumulation under abiotic stresses (Qiao *et al*., 2015).

Sequence analysis revealed that annexin may have a post-transcriptional modification site such as a phosphorylation site, which may be a substrate for protein kinases (Jami *et al*., 2012). Evidence obtained using a tandem affinity purification approach to identify protein complexes suggests that annexin may interact with various kinases, including receptor-like kinase, sterile-20 (Ste20)-like kinase, calcium/calmodulin-dependent protein kinase and casein kinase (Rohila *et al*., 2006). In addition to the MAPK cascade, Ca^2+^ signaling is another critical pathway triggered by environmental stimuli and developmental cues. Calcium-dependent protein kinases (CDPKs) are serine/threonine protein kinases that function as one of the best characterized Ca^2+^ sensors in plants. Previous studies suggested that CDPKs are involved in the responses of plants to various abiotic stresses, including salt, drought, hormonal stimuli and oxidative stress (Kobayashi *et al*., 2007; Zhu *et al*., 2007; Asano *et al*., 2012; Rahoui *et al*., 2017). *OsCPK24* (same as OsCDPK24), the rice CDPK gene, increases the tolerance of rice seedlings by reducing the thioltransferase activity of OsGrx10 to increase proline and glutathione contents under cold stress (Liu *et al*., 2018). The transcription factor protein OsDi19-4 interacts with OsCDPK14/OsCPK24 and is further phosphorylated by OsCDPK14/OsCPK24, acting in response to ABA signaling (Wang *et al*., 2016). However, the interactions between protein kinases CDPKs and annexins that are involved in the regulation of antioxidant defenses in ABA signaling remain to be determined.

The aerenchyma is a normal morphological structure utilized by plants to adapt to the waterlogging and submergence of soil, promoting internal oxygen diffusion (Colmer, 2003; Licausi *et al*., 2011; Licausi, 2013). Ultra structural results suggest that cortical cell death occurs constitutively (Joshi and Kumar, 2012) via a process termed constitutive aerenchyma formation. Under oxygen-deficient conditions, many plant roots will induce aerenchyma development to ensure internal oxygen transport from the shoots to roots, thus facilitating gas diffusion (Campbell and Drew, 1983; Colmer, 2003). Aerenchyma formation is regulated through reactive oxygen species (ROS) in plants (Colmer and Pedersen, 2008; Rajhi *et al*., 2011; Yamauchi *et al*., 2014). Hydrogen peroxide (H_2_O_2_), a type of ROS, plays a key role in programmed cell death and lysigenous aerenchyma formation (Yamauchi *et al*., 2017).

In this study, we identified a putative annexin family gene in rice, *OsANN4*. We investigated that *OsANN4* alleviates ABA-induced root aerenchyma formation via modulating H_2_O_2_ accumulation. Moreover, OsANN4, a calcium-binding annexin protein, is an interacting protein of kinase OsCDPK24 and further be phosphorylated by OsCDPK24, and responses to ABA treatment. Thus, we have discovered a proposed mechanism for annexin and protein kinase in H_2_O_2_ production in aerenchyma formation to response ABA.

## Materials and methods

### Vector construction for recombinant protein expression and rice transformation

Total RNA was extracted from 7-day-old seedlings of WT plants. Full-length *OsANN4* cDNA without the stop codon was amplified with primers P1 and P2 via reverse transcription PCR. For sequencing and subcloning, the products were initially ligated into a PMD18-T vector. After sequencing, the fragment was cut with *Xba*I and *Kpn*I and inserted into the PMDC83 expression vector driven by a CaMV35S promoter. To construct the RNA interference vector, the 347-bp coding sequence of *OsANN4* was amplified with primers P3 and P4, digested with *Sac*I and *Spe*I followed by *Bam*HI and *Kpn*I, and subsequently ligated into the pTCK303 vector.

For the Ca^2+^-binding assay, the expression vector pET28a-*OsANN4* was constructed. Primers P5 and P6 were designed, and the full-length cDNA encoding *OsANN4* was subcloned into *E. coli* between the *Eco*RI and *Hin*dIII sites in an orientation such that the cDNA sequence was followed by a His tag.

For the pull-down assay, the expression vector pGEX4T-1-*OsANN4* was constructed. The full-length cDNA encoding OsANN4 was ligated into pGEX4T-1 using primers P5 and P6 to obtain the cDNA sequence followed by a GST tag. The expression vector pET28a-*OsCDPK24* was constructed by using primers P7 and P8.

For the LCI assay, the full-length cDNA encoding *OsANN4* was ligated into pCAMBIA-Cluc by using primers P9 and P10, and *OsCDPK24* was ligated into pCAMBIA-Nluc by using primers P11 and P12.

For the construction of pET28a-*OsANN4(S13A*), the primers P5 and P6 were used and the vector of OsANN4(S13A)-Cluc was constructed by using primers P9 and P10. Site-directed mutagenesis was finished by a fast mutagenesis kit (Tiangen, China).

All primer sequences used in these experiments are listed in Table S1. The expression vectors pET28a-*OsANN4* and pGEX4T-1-*OsANN4* were then transformed into the *E. coli* Rosetta strain. In addition, other constructs were introduced into *Agrobacterium* EHA105 and transformed into rice calli. Transgenic rice plants were generated as previously described.

### Plant materials and stress treatment

The rice (*Oryza sativa* subsp. japonica) cultivar Nipponbare was used in this study and as the WT control in all experiments. Rice seeds were surface-sterilized with 50% NaClO for 20 min, rinsed 10 times with sterile distilled water and then grown on 1/2 MS medium. The rice plants were grown in a standard culture solution in a greenhouse with a light/dark cycle of 16/8 h and 50% relative humidity at 28/25 ºC (day/night). For the ABA sensitivity assay, the seeds were planted on 1/2 MS medium supplemented with 10 µM ABA for 12 days. Control seeds were planted on 1/2 MS medium and cultured with water after 12 days.

### RNA isolation, RT-PCR, and quantitative RT-PCR analysis

Total RNA from different tissues of rice plants was isolated using the TaKaRa RNAiso Plus kit. Purified RNA (2 μg) was incubated with DNase 1 (RNase-Free DNase, Thermo Fisher, USA) at 37 °C for 30 min. First-strand cDNA was synthesized using the PrimeScript™ First Strand cDNA Synthesis Kit (TaKaRa, Japan) to perform RT-PCR. For qRT-PCR, 1μg purified total RNA was used to obtain the first-strand cDNA with the PrimeScript™ RT reagent Kit with gDNA Eraser (TaKaRa, Japan). The Takara SYBR ® Premix Ex Taq™ II was used to perform the qRT-PCR assay using primers P13 and P14. The C1000 Real-Time PCR instrument (Bio-Rad, USA) was used for qRT-PCR. *OsACTIN1* (Os03g0718100) was used as an internal control for the normalization of all data by using Primers P15 and P16 in this experiment. Three independent biological replicates were assayed.

### Sectioning and microscopy of root tips

Adventitious roots were obtained from 1/2 MS medium supplemented with 10 μM ABA. After embedding in resin (SPI, USA), the embedded materials were measured and marked at 3 mm and 5 mm from the root tip. Then the transverse sections of 1 μm thickness were acquired on a RM 2265 rotary microtome (Leica, Germany). And the resin section was stained with toluidine blue, and examined and photoed by a DVM6a 3D microscope (Leica, Germany).

### Measurement of H_2_O_2_ content

H_2_O_2_ content in rice seedlings was determined using the Hydrogen Peroxide Assay Kit (Beyotime, China). According to the instructions, the samples were fully homogenized in lysis buffer. Fifty microliters of supernatant was added to 100 μl of test solution and then placed at 30□ for 30 min. Readings at a wavelength of 560 nm were measured immediately. Then, H_2_O_2_ content was calculated according to the standard curve.

### Superoxide dismutase and catalase activity assays

To determine the total soluble protein content, rice seedlings (0.2 g, fresh weight, FW) were homogenized in 1.0 ml of extraction buffer (50 mM phosphate, 1% (w/v) polyvinyl pyrrolidone, pH 7.8) in a mortar on ice. The plant tissue was ground into a homogenate and then centrifuged at 4 ºC for 20 min at 8000 rpm. The supernatants were used for assaying the protein content and enzyme activities. The total soluble protein contents were then determined by the Bradford method using bovine serum albumin (BSA, Solarbio, China) as the standard. All spectrophotometric analyses were performed on a METASH UV-5200PC spectrophotometer (Shanghai, China).

### Assaying the in situ localization of H_2_O_2_

DAB staining was performed following a published method with some modifications. 7-day-old rice seedlings were treated with 10 μM ABA, and then the leaves were detached and immersed in a solution containing 1 mg/ml DAB prepared in HCl-acidified (pH 3.8) water at 25 ºC for 10 h and the roots were immersed in DAB solution for 4h. The samples were then incubated in a boiling solution (60% ethanol, 20% glycerol and 20% ethylic acid) to wash off green color and then observed and imaged under a DVM6a 3D microscope (Leica, Germany).

### Yeast two-hybrid analysis

The *OsANN4* coding sequences were cloned into the BD (pGBKT7) vector between the *Nde*I and *Eco*RI sites with primers P17 and P18. The *OsCDPK24* (Os11g0171500) coding sequence was cloned into AD (pGADT7) vectors by using primers P19 and P20. The expression vectors pGADT7-*OsCDPK24* and pGBKT7-*OsANN4* were co-transformed into the yeast strain AH109 via the lithium acetate method (Clontech Yeast Protocols and book). Colonies were then transferred to SM-LWHA and SM-LW. A positive control interaction between the 53 protein and SV40 protein and a negative control interaction between the Lam protein and SV40 protein were observed.

### Pull down assays

For induction and purification of OsCDPK24, the plasmid pET28a-*OsCDPK24* was induced at 18□ overnight in *E. coli* strain Rosetta and purified with Ni-NTA Resin. OsANN4-GST and GST were induced at 18□ overnight in *E. coli* strain Rosetta and purified with glutathione Sepharose 4B beads (GE). Supernatant containing GST or OsANN4-GST was incubated with OsCDPK24-His in 0.5 ml of interaction buffer (25 mM Tris pH 7.2, 150 mM NaCl) overnight at 4□. GST beads were added and incubated with proteins for 2 h on a rotating wheel at 4□ followed by washing five times with wash buffer (25 mM Tris pH 7.2, 150 mM NaCl, 0.1% NP-40). Then, the samples were boiled with 120 μl of 1× SDS loading buffer at 100□ for subsequent western blot analysis.

### Luciferase complementation imaging (LCI) assay

*OsANN4* and *OsCDPK24* were constructed by using pCAMBIA-Cluc and pCAMBIA-Nluc, respectively, and were transformed into *N. benthamiana* leaves by *Agrobacterium* strain GV3101. Agrobacterium transformants containing p*OsANN4*-Cluc, p*OsCDPK24*-Nluc, empty control pCAMBIA-Cluc and pCAMBIA-Nluc were adjusted to OD_600_=0.5-0.6 and then injected into tobacco leaves. After spraying the tobacco leaves with 2.5 mM D-luciferin (Goldbio, USA), fluorescent signals were detected and photographed post infiltration by using a Fusion FX7 (Vilber, France) imaging system. The cold luminescence signals values of OsANN4 and OsCDPK24 were calculated by Fusion FX7 (Vilber, France).

### In vitro kinase assays

Briefly, 5 μg of OsANN4-His was incubated with 1.5 μg of protein kinase OsCDPK24-His in kinase buffer (20 mM Tris-HCl (pH 7.5), 10 mM MgCl_2_, 100 mM NaCl, 1 m DTT), and 2 mM ATP was added. The reactions were incubated at 37 °C for 10 h and stopped by boiling with 6× SDS loading buffer. After adding 50 mM Phos-Tag (APExbio, China) and 100 mM MnCl_2_, the samples were separated by SDS-PAGE and then transferred to nitrocellulose membranes. The signals were detected with a His-tag mouse monoclonal antibody (CWbio, China). Mass spectrometry to detect phosphorylation was finished by Applied Protein Technology Co. Ltd company (Shanghai, China).

### Fluorescence measurements of OsANN4

This experiment was finished according to the method described in the study previously (Zhou *et al*., 2011). 2 mM Ca^2+^ was added into 2 μM recombinant protein, and the fluorescence signals was measured by fluorescence spectrophotometer (F-4600; Hitachi, Japan).

### Subcellular localization of OsANN4

To examine the subcellular distribution of OsANN4, the 35S::OsANN4-GFP vector was constructed. The OsANN4 coding region was fused to GFP using the PMDC83 backbone to construct CaMV35S::OsANN4-GFP, and its expression was driven by the 35S promoter. The constructs were introduced into *Agrobacterium* EHA105 and then transformed into rice calli. Transgenic rice plants expressing OsANN4-GFP were generated as previously described. The signal of OsANN4-GFP protein was examined using confocal laser-scanning microscopy (Zeiss LSM710, Germany).

## RESULTS

### OsANN4 responses to exogenous abscisic acid in rice

We cloned and obtained a putative annexin family gene in rice, *OsANN4*, which is a ortholog of *AtANN4* (Fig. 1A). The genomic sequence of *OsANN4* consists of 5 exons and 4 introns, and the open reading frame is predicted to encode a protein with 320 amino acids. Based on sequence searches of the NCBI database, the OsANN4 protein contains three annexin domain architectures, the so-called annexin repeats, which comprise segments of 32, 57, and 67 amino acid residues. Analysis of the 1794-bp promoter sequence upstream of *OsANN4* reveals that the *OsANN4* promoter contains multiple cis-acting elements that may be associated with abiotic stress (Fig. S1). Then we treated 7-d-old seedlings with 10 μM ABA and detected the expression of *OsANN4* at 0 h, 0.5 h, 1 h, 3 h, 6 h, 12 h and 24 h. The results showed that ABA treatment induced the expression of *OsANN4* and the highest expression level after ABA treatment for 6 h, which showed that *OsANN4* may respond to exogenous abscisic acid (Fig. 1B).

**Fig. 1.**
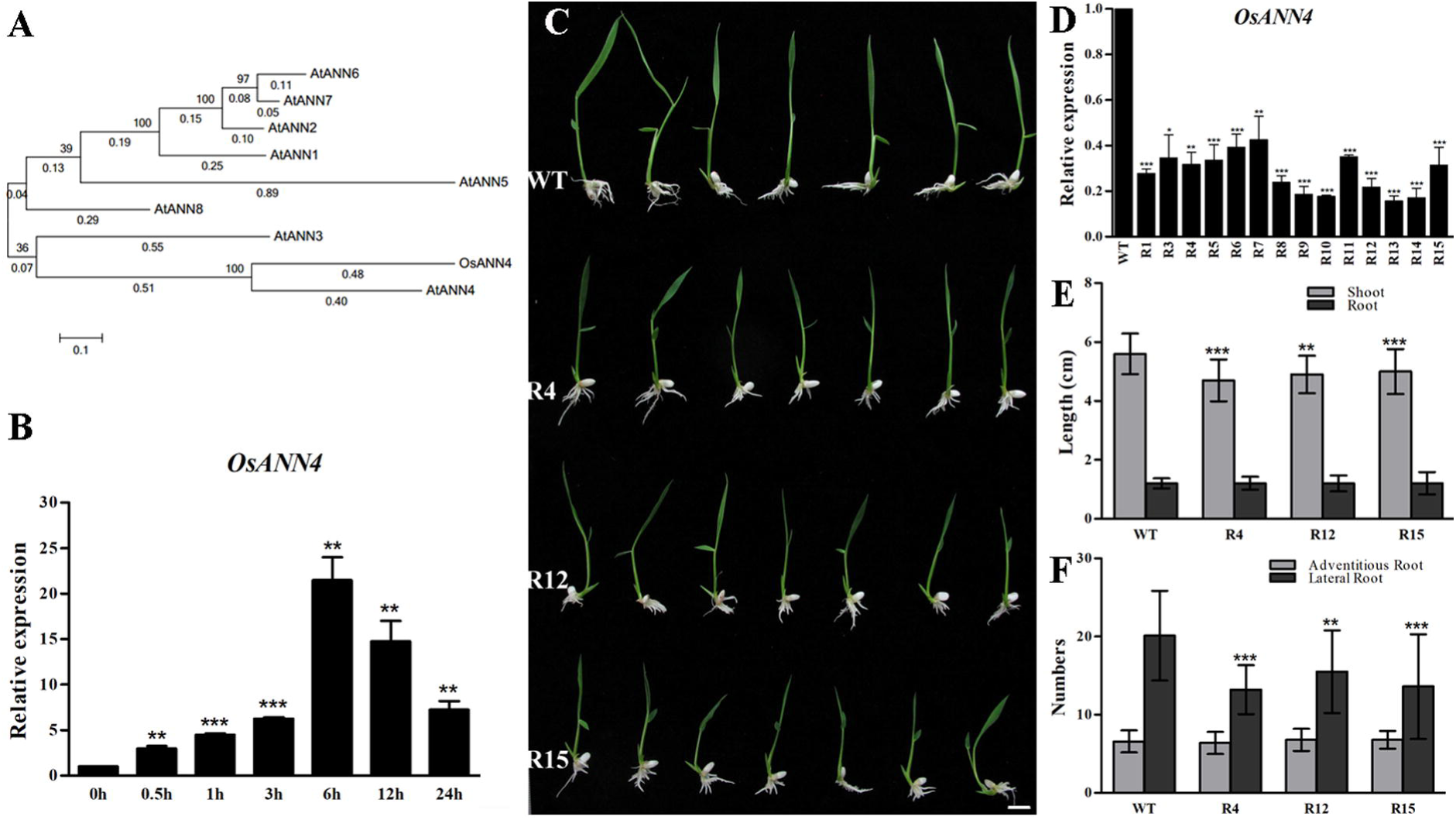
*OsANN4* responses to exogenous abscisic acid in rice. **A** Phylogenic tree of *OsANN4* and the *Arabidopsis* orthologue AtANNs. **B** *OsANN4* transcript expression levels under ABA treatment. **C** WT and *OsANN4* RNAi lines were grown for 12 d in 10 μM ABA medium. The figure shows the representative results of five replicates with T2 generation rice. Scale bars are 1 cm. **D** Relative transcript levels of *OsANN4* in *OsANN4* RNAi transgenic rice. E. The length of shoots and roots in 12-d-old WT and *OsANN4* RNAi plants with 10 μM ABA treatment. **F** The numbers of adventitious roots and lateral roots in 12-d-old WT and *OsANN4* RNAi lines with 10 μM ABA treatment. Values represent the mean ± SD from three independent experiments. Statistical significance was determined by Student’s t-test, *P<0.05, **P<0.01, ***P<0.001.

To test whether *OsANN4* is sensitive to ABA treatment, fifteen independent knocking down OsANN4 transgenic lines were generated. Quantitative real-time PCR analyses showed that *OsANN4* was down regulated in all the RNAi lines (Fig. 1D). The homozygous plants of R4, R12 and R15 showed obviously decreased transcriptional level of *OsANN4* and to be used for the further analyzed. Seeds from a wild-type (WT) line and three RNAi lines (R4, R12 and R15) were planted on 1/2 Murashige and Skoog (MS) medium supplemented with 10 μM ABA, and on 1/2 MS medium as a control. Twelve days later, both WT and RNAi plants showed no significant difference at the stage of early seedling growth without ABA treatment (Supplementary Fig. S2A). Otherwise, the shoots of RNAi lines were shorter than WT with ABA (Fig. 1C, E). Furthermore, the length of the root and the numbers of adventitious roots were not significantly different from that of the wild type after ABA treatment, whereas the number of lateral roots was significantly less relative to wild type line (Fig. 1E, F). This result indicates that the RNAi seedlings are sensitive to ABA in the early seedling stage.

To understand if *OsANN4* expression is involved in ABA response, we further examined the expression levels of *OsANN4* and several genes related ABA biosynthesis and response in RNAi lines and WT treated with 10 μM ABA for 0h, 0.5h, 1h, 3h and 6h. The expression levels of *OsANN4* in RNAi lines were always lower relative to WT (Supplementary Fig. S2B). Under both normal and ABA treatment, *OsNCED4* and *RAB16A* expression were decreased in the RNAi lines, especially in 6h (Supplementary Fig. S2C, S2E). However, the expression of the *OsZEP* transcripts did not differ between the WT and RNAi lines (Supplementary Fig. S2D). These data sufficiently demonstrated that *OsANN4* is closely related to ABA response.

### OsANN4 modulates ROS in response to ABA in rice roots

More evidence suggests that ABA-enhanced stress tolerance is associated with the induction of the antioxidant defense system to protect plant cells against oxidative damage (Xing *et al*., 2008; Ozfidan *et al*., 2012). SOD and CAT may protect plant cells from oxidative damage by eliminating ROS, which are important signaling molecules in the ABA pathway. SOD is a critical enzyme that catalyzes O_2_^.−^ to H_2_O_2_, and CAT is a key enzyme that catalyzes H_2_O_2_ to H_2_O and O_2_ which protects plant cells from oxidative damage by eliminating ROS. To assess the effect of knocking down *OsANN4* on antioxidant defense to ABA response, we detected the activities of SOD and CAT in 7-d-old seedlings without or with 10 μM ABA treatment for 1 h. The RNAi lines showed no significant difference in activities of both SOD and CAT relative to those of the WT without treatment. After ABA treatment for 1h, both the SOD and CAT activities of RNAi lines were decreased relative to WT (Fig. 2A, B).

**Fig. 2.**
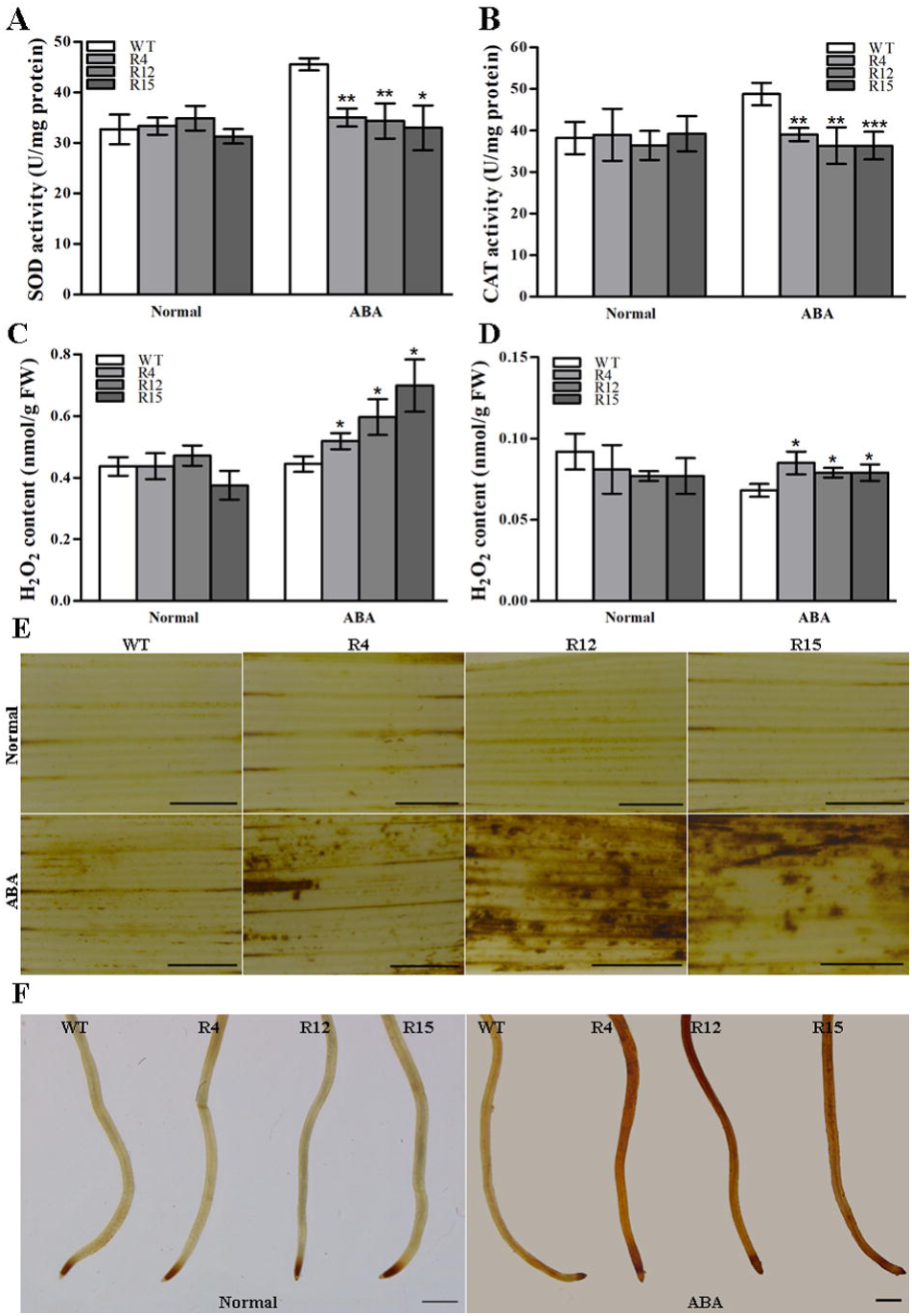
*OsANN4* modulates ROS in response to ABA in rice roots. **A** SOD activities in 7-d-old WT and *OsANN4*-R plants with or without 10 μM ABA treatment for1h. **B** CAT activities in 7-d-old WT and *OsANN4*-R plants with or without 10 μM ABA treatment for 1h. **C** The H_2_O_2_ content in the shoots of 7-d-old WT and *OsANN4*-R plants with or without 10 μM ABA treatment for 1h. **D** The H_2_O_2_ content in the roots of 7-d-old WT and *OsANN4*-R plants with or without 10 μM ABA treatment for 1h. **E** Histochemical detection of H_2_O_2_ in situ in the leaves of WT and *OsANN4*-R plants with or without ABA treatment. **F** Histochemical detection of H_2_O_2_ in situ in the roots of WT and *OsANN4*-R plants with or without ABA treatment. Scale bars for images E. and (f) are 1 mm. Values represent the mean ± SD from three independent experiments. Statistical significance was determined by Student’s t-test, *P<0.05, **P<0.01, ***P<0.001.

We further detected H_2_O_2_ content in 7-d-old RNAi and WT seedlings. The RNAi lines showed increased H_2_O_2_ content in both shoots and roots after 10 μM ABA treatment (Fig. 2C, D). The results were consistent with SOD and CAT activity. Next, we assayed the production of H_2_O_2_ in situ of RNAi lines and WT. Seven-day-old rice seedlings were treated with or without 10 μM ABA for 5h, roots and leaves were then subjected to 3, 3’-diaminobenzidine (DAB) staining respectively. Without ABA treatment, there were no visible differences in rice roots and leaves of WT and RNAi lines. However, increased numbers of darkly stained patches were observed in mesophyll cells and root cells of RNAi lines (Fig. 2E, F), indicating an increased level of H_2_O_2_ in these plants.

### OsANN4 promotes stomatal closure and alleviates H_2_O_2_-induced aerenchyma formation with ABA treatment

It is reported that a certain ROS content resulted in programmed cell death and further promoted aerenchyma formation (Yamauchi *et al*., 2017). The H_2_O_2_ content was obviously increased in the roots of the *OsANN4*-RNAi lines, then we investigate whether *OsANN4* affects aerenchyma formation in adventitious roots. Twelve-day-old *OsANN4-*RNAi lines as well as WT were planted in 1/2 MS medium supplemented with 10 μM ABA. Upon analysis of the transverse sections, aerenchyma formation was observed only in the RNAi lines and not in the WT at 3-mm from root tips.

Furthermore, aerenchyma formation was observed in both RNAi lines and WT at the 5-mm transverse sections (Fig. 3A, B). Then we also detected H_2_O_2_ content of 12-d-old RNAi and WT seedlings with 10 μM ABA treatment. The RNAi lines showed higher H_2_O_2_ content relative to WT with ABA treatment, which may result in an increase in the number of aerenchyma in the RNAi lines (Supplementary Fig. S3).

**Fig. 3.**
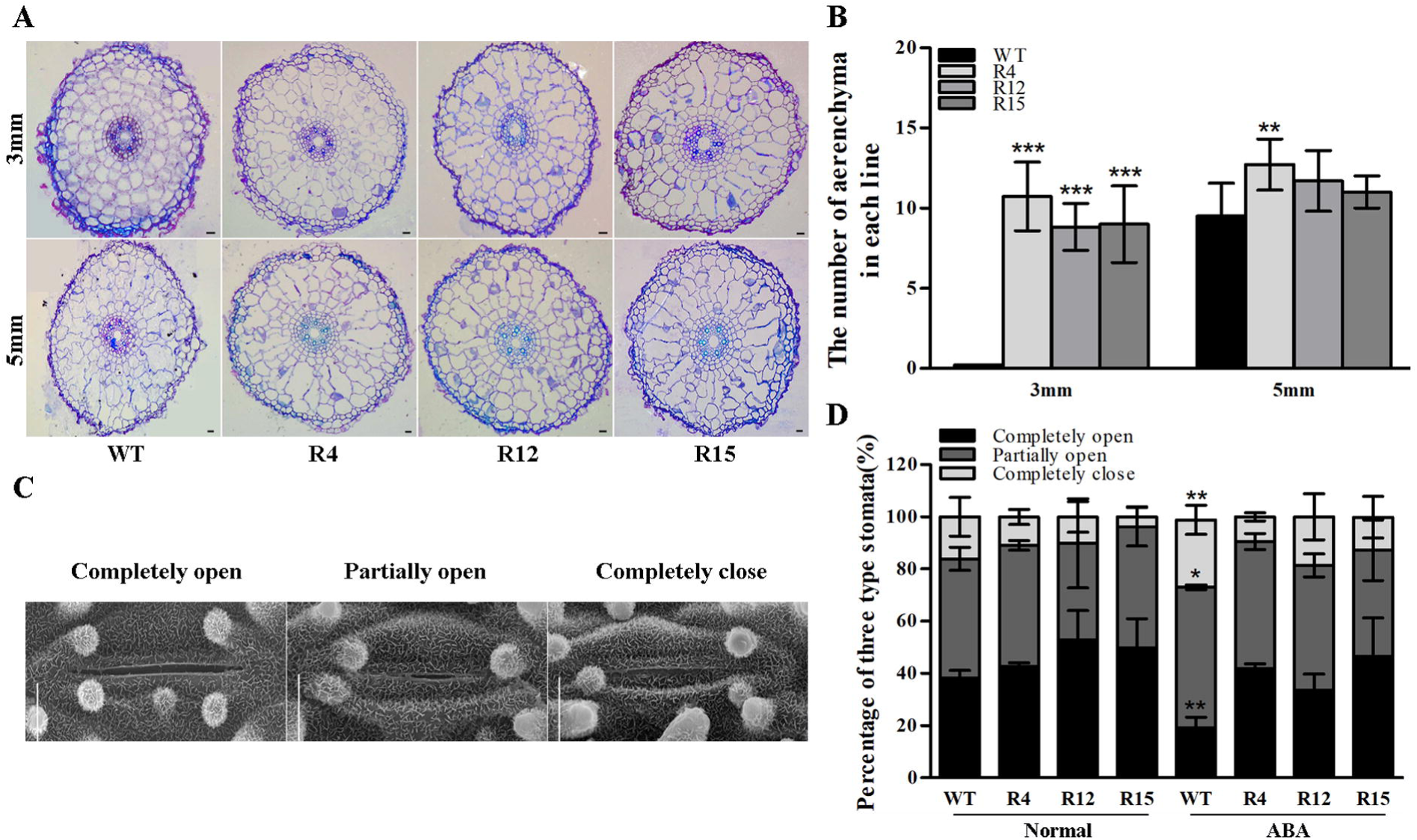
*OsANN4* promotes stomatal closure and alleviates H_2_O_2_-induced aerenchyma formation with ABA treatment. **A** Scanning electron microscopy images of three levels of stomatal apertures. **B** The percentage of three levels of stomatal apertures in the leaves of *OsANN4* RNAi and WT plants under normal and ABA treatment. Values are means ± SD (23≤n≤36). *P<0.05, **P<0.01. **C** Cross sections at 3 mm and 5 mm from the tips of the adventitious roots in WT and *OsANN4*-R plants with 10 μM ABA treatment for 12 d. Sections were stained with toluidine blue. Scale bars are 20 μm. **D** The aerenchyma numbers in the adventitious roots of WT and *OsANN4*-R plants with ABA treatment. Values represent the mean ± SD from three independent experiments. Statistical significance was determined by Student’s t-test, *P<0.05, **P<0.01, ***P<0.001.

We further assessed the stomatal apertures of these plants using scanning electron microscopy images of rice leaves. The proportion of stomatal closure in WT was significantly increased with ABA treatment than that without ABA treatment.

However, for RNAi plants, the statistical showed no significantly differences of stomatal openness and closure with or without 10 µM ABA treatment (Fig. 3C, D). It suggests that *OsANN4* may play a key role in ABA-induced stomatal closure.

### OsANN4 interacts with the protein kinase OsCDPK24

In previous reports, annexins were shown to interact with protein kinases, including SAPKs and CDPKs (Rohila *et al*., 2006; Qiao *et al*., 2015). To further understand how *OsANN4* response to ABA, we used yeast two-hybrid assay to verify several potential rice protein kinase candidates, including Os01g0570500, Os10g0518800, Os01g0869900, Os11g0171500. With the results, we did not find Os01g0570500 and Os10g0518800 interacted with OsANN4 separately. However, Os01g0869900, belongs to sucrose nonfermenting1-related protein kinase2 (SnRK2) family, showed weak interaction with OsANN4. Furthermore, OsCDPK24 (Os11g0171500), a key regulator in response to ABA (Wang *et al*., 2016), showed strong interaction with OsANN4 (Fig. 4A).

**Fig. 4.**
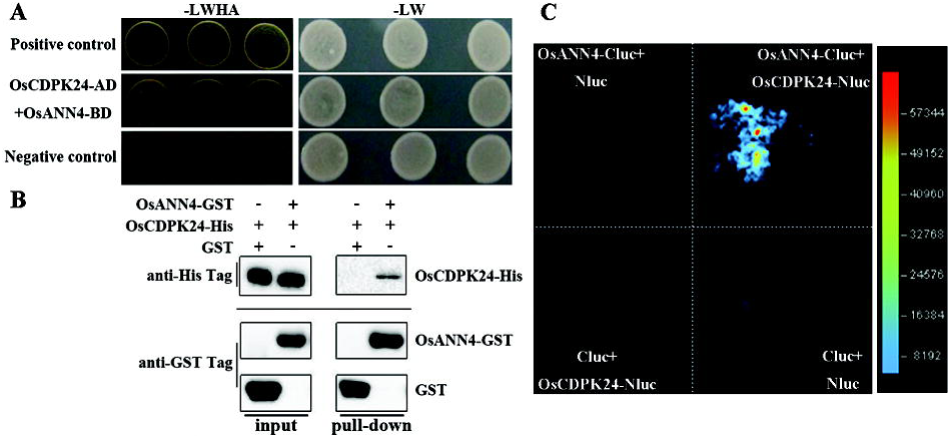
OsANN4 can interact with the protein kinase OsCDPK24. **A** Yeast two-hybrid system was used to detect the interaction between OsANN4 and OsCDPK24. **B** A GST pull-down assay was used to detect the interaction between OsANN4 and OsCDPK24. **C** Firefly luciferase complementation imaging (LCI) assay detecting the interaction between OsANN4 and OsCDPK24. The colored scale bar indicates the luminescence intensity.

To further verify the interactions between OsANN4 and OsCDPK24, the pull-down system in vitro was carried out. OsCDPK24-His and OsANN4-GST recombinant proteins were induced in *Escherichia coli* and purified to perform the pull-down assay. OsANN4-GST could pull down OsCDPK24-His, whereas GST could not, it was further proved the interaction between OsANN4 and OsCDPK24 (Fig. 4B).

We also obtained additional confirmation of the interaction between OsANN4 and OsCDPK24 in *Nicotianana benthamiana* leaves by using a luciferase complementation imaging (LCI) assay. After spraying D-luciferin to tobacco leaves, a fluorescent signal occurred when p*OsANN4*-Cluc and p*OsCDPK24*-Nluc were present simultaneously, which showed that OsANN4 can interact with OsCDPK24 (Fig. 4C).

### The 13th serine residue is a key phosphorylation site for OsANN4

To understand the mechanism underlying the interaction of OsANN4 and OsCDPK24, we performed a phosphorylation assay in vitro to determine whether OsANN4 is a substrate of OsCDPK24. The Phos-tag reagent was used to separate phosphorylated proteins from non-phosphorylated proteins according to their different migration rates. When purified OsANN4-His and OsCDPK24-His were incubated together, phosphorylated OsANN4 bands were detectable with a His-tag antibody. The phosphorylation level of OsANN4 increased slightly after the addition of 5 μM and 500μM Ca^2+^. These results indicated that OsANN4 can be phosphorylated by OsCDPK24, especially in the presence of Ca^2+^ (Fig. 5A).

**Fig. 5.**
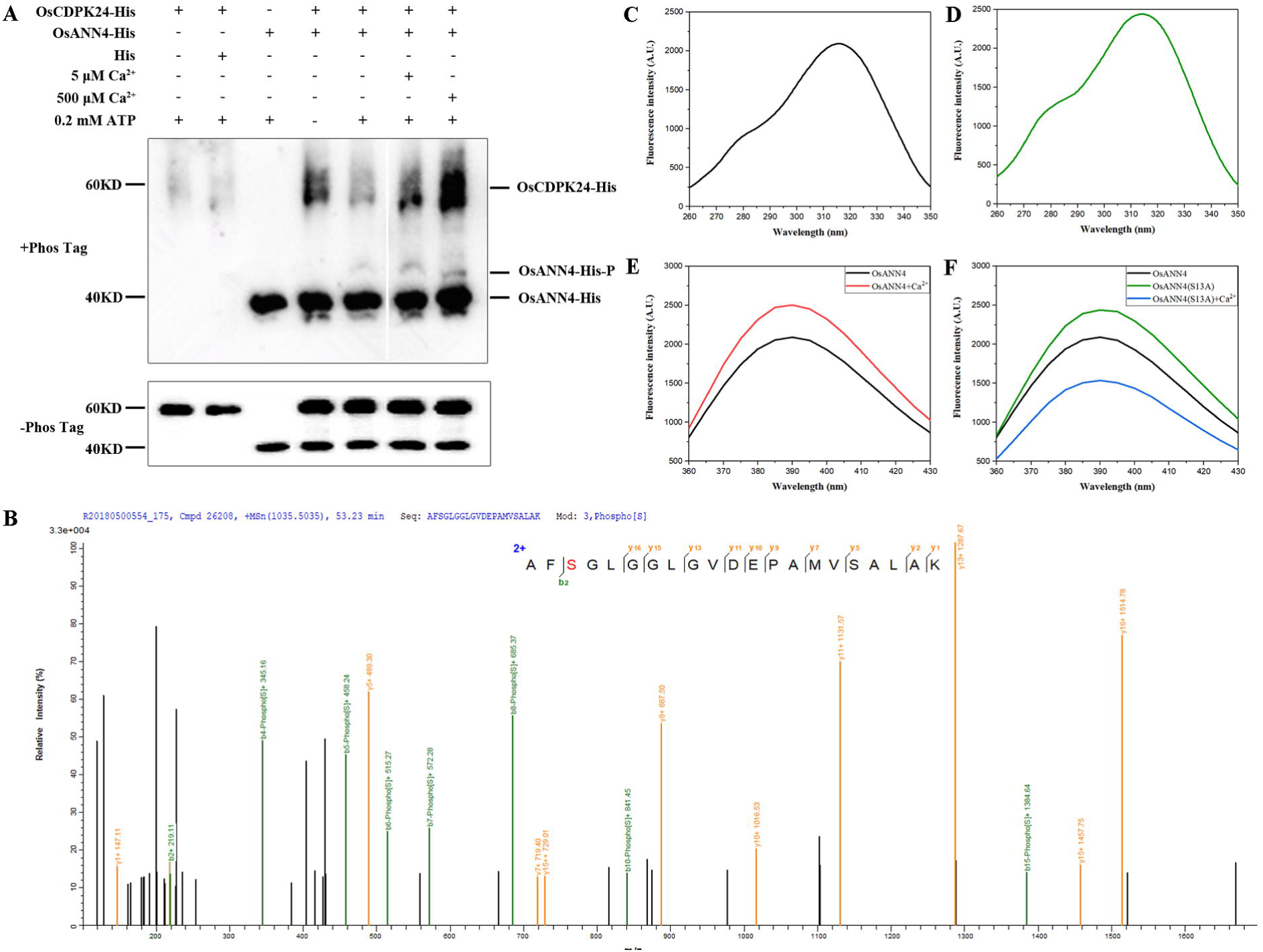
OsANN4 is a Ca^2+^-binding protein that can be phosphorylated by OsCDPK24. **A** An in vitro kinase assay was performed with purified OsANN4-His and OsCDPK24-His using a Phos-tag gel. Signals were detected by using a His-tag antibody. **B** Mass spectrometry analysis of OsANN4 phosphorylation. **C** The fluorescence excitation of the purified OsANN4-His protein. **D** The fluorescence excitation of the purified OsANN4(S13A)-His protein. **E** The emission spectra of the purified OsANN4-His protein. **F** The emission spectra of the purified OsANN4(S13A)-His protein.

To further demonstrate that OsANN4 was a substrate of OsCDPK24, mass spectrometry was performed to detect specific phosphorylation sites of OsANN4 when OsCDPK24 was present. Mass spectrometry analysis showed that OsANN4 can be phosphorylated by OsCDPK24, and the OsANN4 phosphorylation site was the 13th amino acid, which is a serine (Fig. 5B).

### The interaction of OsANN4 and OsCDPK24 plays a role in the response to ABA

To determine whether the interaction of OsANN4 and OsCDPK24 plays a role in the response to ABA, we performed 10 μM ABA treatment on tobacco leaves injected with *Agrobacterium* GV3101 containing p*OsANN4*-Cluc and p*OsCDPK24*-Nluc. After ABA treatment, LUC cold luminescence signals increased significantly, which demonstrated the interaction between OsANN4 and OsCDPK24 was enhanced with ABA treatment (Fig. 6A and B). These results indicated that OsANN4 interacted with OsCDPK24, and this interaction increased under ABA treatment, thus, the OsANN4-OsCPK24 interaction may play a role in the response to ABA in rice.

**Fig. 6.**
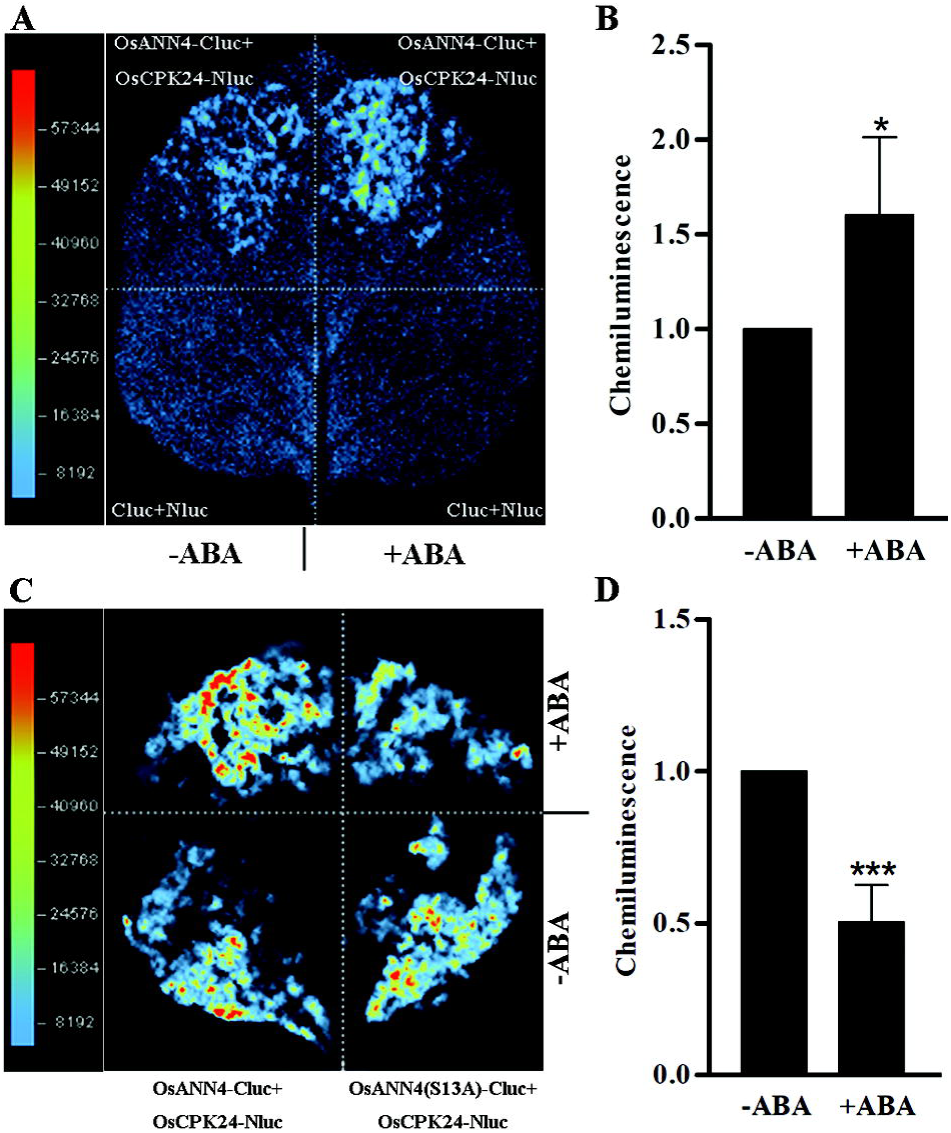
The interaction between OsANN4 and OsCDPK24 involved in response to ABA signals. **A** Firefly luciferase (LUC) complementation imaging(LCI) assay detecting the interaction between OsANN4 and OsCDPK24 with or without ABA treatment. The colored scale bar indicates the luminescence intensity. **B** Quantification of the relative cold luminescence signals displayed in the two group of A.. Values represent mean ± SD and the single asterisk represent a significant difference determined by the Student’s test, *P<0.05, n>13. **C** Firefly luciferase complementation imaging (LCI) assay detecting the interaction between OsANN4-(S13A) and OsCDPK24 with ABA treatment. The colored scale bar indicates the luminescence intensity. **D** Quantification of the relative cold luminescence signals displayed between OsANN4-(S13A) and OsCDPK24 with ABA treatment. Values represent the mean ± SD and statistical significance was determined by Student’s t-test, ***P<0.001, n>12.

To demonstrate whether the phosphorylation of OsANN4 plays a role in response to ABA signaling, we mutated the serine (S) residue with non-phosphorylatable alanine (A) to inhibit the phosphorylation of OsANN4 and constructed the p*OsANN4 (S13A*)-Cluc vector to perform an LCI assay. We found that after mutation of phosphorylation site in OsANN4, OsANN4 still can interact with OsCDPK24. After spraying with ABA, the responses of OsANN4 (S13A)-Cluc and OsCDPK24-Nluc to ABA were significantly weakened (Fig. 6C, D). This suggests that this key phosphorylation site plays an important role in *OsANN4* response to ABA.

### OsANN4 is a Ca^2+^-binding protein that locates to the cell periphery

Previous studies have shown that the subcellular localization of annexin may be altered due to environmental stimuli (Baucher *et al*., 2012; Qiao *et al*., 2015). To explore whether OsANN4 altered its subcellular localization with ABA treatment, 35Spro::OsANN4-GFP was introduced into *Agrobacterium* EHA105 and then transformed the rice callus. Twenty-five independent transgenic lines were obtained, and 3 lines were used for further analyses. We observed OsANN4 was located to the cell periphery by a confocal laser-scanning microscope (Zeiss LSM710). After 10 μM ABA treatment, the signal of OsANN4-GFP remained unchanged (Supplementary Fig. S4).

Annexins are considered to be a class of protein interacting with biological membranes in a calcium dependent or independent manner. In this study, we detected the Ca^2+^ binding activity of OsANN4. The fluorescence level of OsANN4-His recombinant protein was determined by fluorescence spectrophotometer (F-4600; Hitachi, Japan). Upon excitation at 315 nm (Fig. 5C), the fluorescence emission spectrum showed the maximum fluorescence wavelength (λ max) at 390 nm, where the fluorescence intensity reached approximately 2000 A.U.. The maximum fluorescence intensity of the OsANN4-His recombinant protein was measured again after the addition of 2 mM Ca^2+^. The results showed that the maximum fluorescence wavelength remained unchanged, whereas the fluorescence intensity changed (Fig. 5E), indicating that the OsANN4 has Ca^2+^ binding activity and may further change the conformation of the protein.

Since the phosphorylation site of OsANN4 plays a role in the ABA response, we examined whether this phosphorylation site affects its binding to Ca^2+^. OsANN4 (S13A)-His recombinant protein was induced and purified to perform the above fluorescence assay. The results showed that upon excitation at 315 nm (Fig. 5D), the fluorescence emission spectrum of OsANN4 (S13A)-His recombinant protein remained unchanged whereas the fluorescence level has changed (Fig. 5F), which indicated that the phosphorylation site of OsANN4 does not affect its binding to Ca^2+^ but may change the conformation of OsANN4.

## Discussion

### OsANN4 modulates ROS production to alleviate aerenchyma formation

Recent results have revealed that annexins are involved in the responses to various abiotic stresses including salt, drought, oxidative and other stresses (Clark *et al*., 1998; Lee *et al*., 2004; Clark *et al*., 2012; Jami *et al*., 2012; Richards *et al*., 2014; Qiao *et al*., 2015). Evidence has shown that induction of the antioxidant defense system protects plant cells against oxidative damage (Mittler *et al*., 2011; Richards *et al*., 2014; Del Rio, 2015; Qiao *et al*., 2015). And ROS are important signaling molecules, especially in the ABA signaling pathway. Herein, we reported *OsANN4*, a ortholog of *AtANN4*, modulates H_2_O_2_ accumulation to alleviate root aerenchyma formation with exogenous abscisic acid. Our results showed that knocking down *OsANN4* decreased the activity of SOD and CAT with ABA treatment, and more H_2_O_2_ content was measured (Fig. 2), further resulted in earlier aerenchyma formation (Fig. 3A, B). We conclude that *OsANN4* promotes redox reactions, potentially regulating SOD and CAT to scavenge H_2_O_2_.

Hydrogen peroxide (H_2_O_2_), is also an essential secondary messenger to be involved in ABA-induced stomatal closure. Interestingly, in our study, a high level of H_2_O_2_ content did not result in a high proportion of stomatal closure in *OsANN4*-RNAi lines (Fig. 3D). Abscisic acid causes the Ca^2+^ concentration in cytoplasmic to elevate in guard cells via Ca^2+^ influx, results in stomatal closure (Hedrich, 2016; Kong *et al*., 2016). We speculate that OsANN4, a putative Ca^2+^ channel, may function in stomatal closure by affecting Ca^2+^ influx under ABA treatment.

### OsANN4 mediates Ca^2+^ and interacts with OsCDPK24 in response to ABA

In rice, a previous study showed that a rice calcium-dependent protein kinase *OsCDPK12* can induce expression of the antioxidant genes *OsAPX2* and *OsAPX8* under salt stress and reduce the salt-induced accumulation of H_2_O_2_ (Asano *et al*., 2012), suggesting that *OsCDPK12* positively regulates ROS detoxification by controlling the expression of antioxidant genes. *ZmCPK11* has been proven to be involved in ABA-induced antioxidant defense and to function upstream of *ZmMPK5* in ABA signaling in maize (Ma *et al*., 2016). However, the regulatory network regulating antioxidant defense in ABA signaling remains to be elucidated. In this study, we demonstrated that OsANN4 interacts with the kinase OsCDPK24 (Fig.4), and furthermore, this interaction plays a role in the ABA response (Fig. 6). Moreover, OsANN4 is a substrate for OsCDPK24 and OsANN4 phosphorylation site is the 13th serine (Fig. 5), which is a key site for phosphorylation. After mutated the serine residue with non-phosphorylatable alanine (A) to inhibit the phosphorylation of OsANN4, the conformation of OsANN4 has changed (Fig. 5F) and the interaction of OsANN4 and OsCDPK24 was weakened (Fig. 6C, D), indicating that under ABA treatment, OsCDPK24 may be more inclined to interact with the phosphorylated OsANN4, rather than the non-phosphorylated state. RCAR is a component of the ABA receptor complex, and PP2C phosphatases inhibit the activity of some kinases, similar to OsCDPK, which can be dephosphorylated and promote ion channel expression (Kudla *et al*., 2018) in response to ABA. However, whether CDPKs are involved in ABA-induced antioxidant defense remains to be determined.

Annexins are traditionally perceived as Ca^2+^-dependent phospholipid-binding proteins, which usually contain a characteristic type II Ca^2+^-binding residue in each corresponding repeat in vertebrates. In plant annexins, the type II Ca^2+^-binding residues are absent in repeats 2 and 3, and no type II Ca^2+^-binding residues are found in OsANN4 (predicted at ScanProsite http://ca.expasy.org/tools/scanprosite/).

However, our results indicate that OsANN4 has the Ca^2+^ binding activity, which could be perceived as evidence that annexins can also function in their Ca^2+^-free conformation in an unknown intricate fashion, thus increasing the functional diversity of annexins. After mutating the phosphorylation site of OsANN4, the protein conformation of OsANN4 is altered, but it still can bind to Ca^2+^ (Fig. 5F). What’s more, some data have indicated that ion channels or transporters act as targets of Ca^2+^ signaling networks (Whalley *et al*., 2011; Edel and Kudla, 2015; Kudla *et al*., 2018). As a putative ion channel, OsANN4 can interact with OsCDPK24, supporting the existence of potential connection of the annexin, calcium and protein kinase in response to stress.

### A possible modle for OsANN4 to respond to ABA

The conformation of PYL can be changed after binding ABA, thus enhancing the stability of the complexes of PYL and PP2C, resulting in a reduced inhibitory effect of PP2C on kinases (SnRK or CDPK, etc.), which then activates transcription factors, ion channels or other ABA-responsive genes (Finkelstein, 2013; Kudla *et al*., 2018; Wang *et al*., 2018). Based on our experimental results, we proposed a working model for OsANN4 functioning as a key protein in modulating H_2_O_2_ accumulation to alleviate root aerenchyma formation with exogenous abscisic acid (Fig. 7). OsANN4 is a Ca^2+^-binding protein, and it can interact with and be phosphorylated by OsCDPK24 at the 13th serine. When ABA is present, PP2C binds to the PYL receptor, may result in a decrease in the inhibitory effect of PP2C on OsCDPK24. The interaction between OsANN4 and OsCDPK24 is enhanced to response ABA. When OsCDPK24 interacts with OsANN4, it may be more inclined to phosphorylate OsANN4 under ABA treatment. *OsANN4* promotes O_2_^.−^ to H_2_O_2_ by increasing SOD activity, then converts H_2_O_2_ into H_2_O through CAT activity, thereby reducing ROS content (O_2_^.-^ and H_2_O_2_). As a consequence, *OsANN4* can further alleviate programmed cell death and inhibit the formation of aerenchyma.

**Fig. 7.**
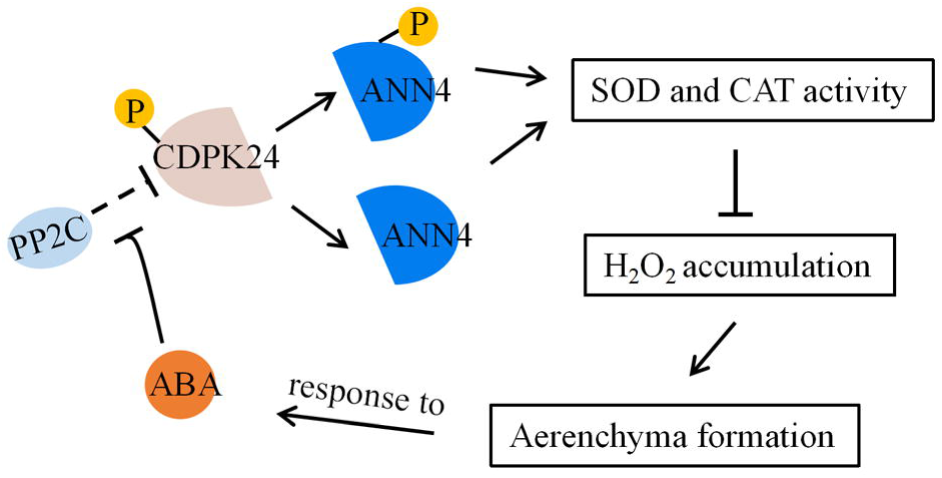
A proposed model for OsANN4 in response to ABA. The conformation of PYL can be changed after binding ABA, thus enhancing the stability of the complexes of PYL and PP2C, resulting in reducing the inhibitory effect of PP2C on OsCDPK24. OsCDPK24 interacts with OsANN4 and further phosphorylates OsANN4. OsANN4 enhances SOD and CAT activities to scavenge H_2_O_2_. As a consequence, OsANN4 can further alleviate the formation of aerenchyma. Under ABA treatment, the interaction between OsANN4 and OsCDPK24 is strengthened, and OsANN4 may be more inclined to be phosphorylated by OsCDPK24.

## Supplementary data

Supplementary data are available at JXB online.

Fig. S1. Analysis of cis-acting elements in the promoter of *OsANN4*.

Fig. S2. The relative transcript levels of *OsANN4*, *OsZEP*, *OsNCED4* and *OsRAB16A* in WT and *OsANN4*-R plants with ABA treatment

Fig. S3. The H_2_O_2_ content in the roots of 12-d-old RNAi and WT seedlings planted in 1/2 MS medium supplemented with or without 10 μM ABA.

Fig. S4. Subcellular localization of OsANN4-GFP in the 3-day-old root tips cell with or without ABA treatmentl.

Table S1. Primer sequences used in plasmid construction and qPCR.

## Acknowledgements

This work was supported by the National Natural Science Foundation of China (31571638, 31340046), Research Fund for the Doctoral Program of Higher Education (20131303110005) and the Advanced Postdoctoral Science Programs Foundation of He’bei Educational Committee (B2016003012).

We thank Dr. Kang Chong of the Institute of Botany, Chinese Academy of Science, for providing the pTCK303 vector. We thank Shanghai Applied Protein Technology Co. Ltd for the technical assistance in LC/MS proteomic assays.

## References

Asano T, Hayashi N, Kobayashi M, et al. 2012. A rice calcium-dependent protein kinase OsCDPK12 oppositely modulates salt-stress tolerance and blast disease resistance. Plant J 69, 26–36

Baucher, M, Perez-Morga, D, and El Jaziri, M. 2012. Insight into plant annexin function, from shoot to root signaling. Plant signaling & behavior 7, 524–528.

Bueso E, Rodriguez L, Lorenzo-Orts L, Gonzalez-Guzman M, Sayas E, Munoz-Bertomeu J, Ibanez C, Serrano R, Rodriguez PL. 2014. The single-subunit RING-type E3 ubiquitin ligase RSL1 targets PYL4 and PYR1 ABA receptors in plasma membrane to modulate abscisic acid signaling. Plant J 80, 1057–1071

Campbell R, Drew MC. 1983. Electron microscopy ofgas space (aerenchyma) formation in adventitious roots of Zea mays L. subjected to oxygen shortage. Planta 157, 350–357

Clark GB, Dauwalder M, Roux SJ. 1998. Immunological and biochemical evidence for nuclear localization of annexin in peas. Plant Physiol Biochem 36, 621–627

Clark GB, Morgan RO, Fernandez MP, Roux SJ. 2012. Evolutionary adaptation of plant annexins has diversified their molecular structures, interactions and functional roles. New Phytol 196, 695–712

Colmer TD. 2003. Aerenchyma and an inducible barrier to radial oxygen loss facilitate root aeration in upland, paddy and deep-water rice (Oryza sativa L.). Ann Bot 91 Spec No, 301–309

Colmer TD, Pedersen O. 2008. Oxygen dynamics in submerged rice (Oryza sativa). New Phytol. 178, 326–334.

Cutler SR, Rodriguez PL, Finkelstein RR, Abrams SR. 2010. Abscisic acid: emergence of a core signaling network. Annu Rev Plant Biol 61, 651–679

Del Rio LA. 2015. ROS and RNS in plant physiology: an overview. J Exp Bot 66, 2827–2837

Ding Y, Cao J, Ni L, Zhu Y, Zhang A, Tan M, Jiang M. 2013. ZmCPK11 is involved in abscisic acid-induced antioxidant defence and functions upstream of ZmMPK5 in abscisic acid signalling in maize. J Exp Bot 64, 871–884

Edel KH, Kudla J. 2015. Increasing complexity and versatility: how the calcium signaling toolkit was shaped during plant land colonization. Cell Calcium 57: 231–246

Finkelstein,R. 2013. Abscisic acid synthesis and response. The Arabidopsis Book 11, e0166–e0166

Fujii H, Chinnusamy V, Rodrigues A, Rubio S, Antoni R, Park SY, Cutler SR, Sheen J, Rodriguez PL, Zhu JK. 2009. In vitro reconstitution of an abscisic acid signalling pathway. Nature 462, 660–664

Giraudat J, Parcy F, Bertauche N, Gosti F, Leung J, Morris PC, Bouvier-Durand M, Vartanian N. 1994. Current advances in abscisic acid action and signalling. Plant Mol Biol 26, 1557–1577

Hedrich R. 2012. Ion channels in plants. Physiol Rev 92, 1777–1811

He Y, Hao Q, Li W, Yan C, Yan N, Yin P. 2014. Identification and characterization of ABA receptors in Oryza sativa. PLoS One 9, e95246

Himmelbach A, Yang Y, Grill E. 2003. Relay and control of abscisic acid signaling. Curr Opin Plant Biol 6, 470–479

Jami SK, Clark GB, Ayele BT, Roux SJ, Kirti PB. 2012. Identification and characterization of annexin gene family in rice. Plant Cell Rep 31, 813–825

Jiang M, Zhang J. 2002. Water stress-induced abscisic acid accumulation triggers the increased generation of reactive oxygen species and up-regulates the activities of antioxidant enzymes in maize leaves. J Exp Bot 53, 2401–2410

Joshi R, Kumar P. 2012. Lysigenous aerenchyma formation involves non-apoptotic programmed cell death in rice (Oryza sativa L.) roots. Physiol Mol Biol Plants 18, 1–9

Klingler JP, Batelli G, Zhu JK. 2010. ABA receptors: the START of a new paradigm in phytohormone signalling. J Exp Bot 61, 3199–3210

Kobayashi M, Ohura I, Kawakita K, Yokota N, Fujiwara M, Shimamoto K, Doke N, Yoshioka H. 2007. Calcium-dependent protein kinases regulate the production of reactive oxygen species by potato NADPH oxidase. Plant Cell 19, 1065–1080

Kong DD, Hu HC, Okuma E, et al. 2016. L-Met activates Arabidopsis GLR Ca 2+ channels upstream of ROS production and regulates stomatal movement. Cell Reports 17, 2553–2561

Kovacs I, Ayaydin F, Oberschall A, Ipacs I, Bottka S, Pongor S, Dudits D, Toth EC. 1998. Immunolocalization of a novel annexin-like protein encoded by a stress and abscisic acid responsive gene in alfalfa. Plant J 15, 185–197

Kudla J, Becker D, Grill E, Hedrich R, Hippler M, Kummer U, Parniske M, Romeis T, Schumacher K. 2018. Advances and current challenges in calcium signaling. New Phytol 218, 414–431

Laohavisit A, Brown AT, Cicuta P, Davies JM. 2010. Annexins: components of the calcium and reactive oxygen signaling network. Plant Physiol 152, 1824–1829

Lee S, Lee EJ, Yang EJ, Lee JE, Park AR, Song WH, Park OK. 2004. Proteomic identification of annexins, calcium-dependent membrane binding proteins that mediate osmotic stress and abscisic acid signal transduction in Arabidopsis. Plant Cell 16, 1378–1391

Licausi F. 2013. Molecular elements of low-oxygen signaling in plants. Physiol Plant 148, 1–8

Licausi F, Kosmacz M, Weits DA, Giuntoli B, Giorgi FM, Voesenek LA, Perata P, van Dongen JT. 2011. Oxygen sensing in plants is mediated by an N-end rule pathway for protein destabilization. Nature 479, 419–422

Liu J, Zhang C, Wei C, Liu X, Wang M, Yu F, Xie Q, Tu Jumin. 2016. The ring finger ubiquitin E3 ligase OsHTAS enhances heat tolerance by promoting H_2_O_2_-induced stomatal closure in rice. Plant Physiology, 170(1), 429

Liu Y, Xu C, Zhu Y, Zhang L, Chen T, Zhou F, Chen H, Lin Y. 2018. The calcium-dependent kinase OsCPK24 functions in cold stress responses in rice. J Integr Plant Biol 60, 173–188

Ma F, Ni L, Liu L, Li X, Zhang H, Zhang A, Tan M, Jiang M. 2016. ZmABA2, an interacting protein of ZmMPK5, is involved in abscisic acid biosynthesis and functions. Plant Biotechnol J 14, 771–782

Ma Y, Szostkiewicz I, Korte A, Moes D, Yang Y, Christmann A, Grill E. 2009. Regulators of PP2C phosphatase activity function as abscisic acid sensors. Science 324, 1064–1068

McCourt P, Creelman R. 2008. The ABA receptors -- we report you decide. Curr Opin Plant Biol 11, 474–478

Melcher K, Zhou XE, Xu HE. 2010. Thirsty plants and beyond: structural mechanisms of abscisic acid perception and signaling. Curr Opin Struct Biol 20, 722–729

Miao Y, Lv D, Wang P, Wang XC, Chen J, Miao C, Song CP. 2006. An Arabidopsis glutathione peroxidase functions as both a redox transducer and a scavenger in abscisic acid and drought stress responses. Plant Cell 18, 2749–2766

Miller G, Suzuki N, Ciftci-Yilmaz S, Mittler R. 2010. Reactive oxygen species homeostasis and signalling during drought and salinity stresses. Plant Cell Environ 33, 453–467

Mittler R, Vanderauwera S, Suzuki N, Miller G, Tognetti VB, Vandepoele K, Gollery M, Shulaev V, Van Breusegem F. 2011. ROS signaling: the new wave? Trends Plant Sci 16, 300–309

Moss SE, Morgan RO. 2004. The annexins. Genome Biol 5, 219

Neill S, Barros R, Bright J, Desikan R, Hancock J, Harrison J, Morris P, Ribeiro D, Wilson I. 2008. Nitric oxide, stomatal closure, and abiotic stress. J Exp Bot 59, 165–176

Ng LM, Melcher K, Teh BT, Xu HE. 2014. Abscisic acid perception and signaling: structural mechanisms and applications. Acta Pharmacol Sin 35, 567–584

Nishimura N, Sarkeshik A, Nito K, et al. 2010. PYR/PYL/RCAR family members are major in-vivo ABI1 protein phosphatase 2C-interacting proteins in Arabidopsis. Plant J 61, 290–299

Ni L, Fu X, Zhang H, et al. 2018. Abscisic acid inhibits rice protein phosphatase PP45 via H_2_O_2_ and relieves repression of the Ca2+/CaM-dependent protein kinase DMI3. Plant Cell doi:10.1105/tpc.18.00506

Ozfidan C, Turkan I, Sekmen AH, Seckin B. 2012. Abscisic acid-regulated responses of aba2-1 under osmotic stress: the abscisic acid-inducible antioxidant defence system and reactive oxygen species production. Plant Biol (Stuttg). 14, 337–346

Park SY, Fung P, Nishimura N, et al. 2009. Abscisic acid inhibits type 2C protein phosphatases via the PYR/PYL family of START proteins. Science 324, 1068–1071

Pekic S, Stikic R, Tomljanovic L, Andjelkovic V, Ivanovic M, Quarrie SA. 1995. Characterization of maize lines differing in leaf abscisic acid content in the field .1. abscisic acid physiology. Ann Bot 75, 67–73

Qiao B, Zhang Q, Liu D, et al. 2015. A calcium-binding protein, rice annexin OsANN1, enhances heat stress tolerance by modulating the production of H_2_O_2_. J Exp Bot 66, 5853–5866

Qu L, Wu C, Zhang F, Wu Y, Fang C, Jin C, Liu X, Luo J. 2016. Rice putative methyltransferase gene OsTSD2 is required for root development involving pectin modification. J Exp Bot 67, 5349–5362

Rajhi I, Yamauchi T, Takahashi H, et al. 2011. Identification of genes expressed in maize root cortical cells during lysigenous aerenchyma formation using laser microdissection and microarray analyses. New Phytol 190, 351–368.

Rahoui S, Martinez Y, Sakouhi L, Ben C, Rickauer M, El Ferjani E, Gentzbittel L, Chaoui A. 2017. Cadmium-induced changes in antioxidative systems and differentiation in roots of contrasted Medicago truncatula lines. Protoplasma 254, 473–489

Richards SL, Laohavisit A, Mortimer JC, Shabala L, Swarbreck SM, Shabala S, Davies JM. 2014. Annexin 1 regulates the H_2_O_2_-induced calcium signature in Arabidopsis thaliana roots. Plant J 77, 36–145

Rohila JS, Chen M, Chen S, et al. 2006. Protein-protein interactions of tandem affinity purification-tagged protein kinases in rice. Plant J 46, 1–13

Santiago J, Rodrigues A, Saez A, Rubio S, Antoni R, Dupeux F, Park SY, Marquez JA, Cutler SR, Rodriguez PL. 2009. Modulation of drought resistance by the abscisic acid receptor PYL5 through inhibition of clade A PP2Cs. Plant J 60, 575–588

Shiono K, Ogawa S, Yamazaki S, Isoda H, Fujimura T, Nakazono M, Colmer TD. 2011. Contrasting dynamics of radial O2-loss barrier induction and aerenchyma formation in rice roots of two lengths. Ann Bot 107, 89–99

Umezawa T, Nakashima K, Miyakawa T, Kuromori T, Tanokura M, Shinozaki K, Yamaguchi-Shinozaki K. 2010. Molecular basis of the core regulatory network in ABA responses: sensing, signaling and transport. Plant Cell Physiol 51, 1821–1839

Wang L, Yu C, Xu S, Zhu Y, Huang W. 2016. OsDi19-4 acts downstream of OsCDPK14 to positively regulate ABA response in rice. Plant Cell Environ 39, 2740–2753

Wang P, Song CP. 2008. Guard-cell signalling for hydrogen peroxide and abscisic acid. New Phytol 178, 703–718

Wang P, Zhao Y, Li Z, et al. 2018. Reciprocal Regulation of the TOR Kinase and ABA Receptor Balances Plant Growth and Stress Response. Molecular Cell 69(1), 100–112

Whalley HJ, Sargeant AW, Steele JF, Lacoere T, Lamb R, Saunders NJ, Knight H, Knight MR. 2011. Transcriptomic analysis reveals calcium regulation of specific promoter motifs in Arabidopsis. Plant Cell 23, 4079–4095

Xing Y, Jia W, Zhang J. 2008. AtMKK1 mediates ABA-induced CAT1 expression and H_2_O_2_ production via AtMPK6-coupled signaling in Arabidopsis. Plant J 54, 440–451

Yamauchi T, Watanabe K, Fukazawa A, Mori H, Abe F, Kawaguchi K, Oyanagi A, Nakazono M. 2014. Ethylene and reactive oxygen species are involved in root aerenchyma formation and adaptation of wheat seedlings to oxygen-deficient conditions. J Exp Bot 65, 261–273.

Yamauchi T, Yoshioka M, Fukazawa A, Mori H, Nishizawa NK, Tsutsumi N, Yoshioka H, Nakazono M. 2017. An NADPH oxidase RBOH functions in rice roots during lysigenous aerenchyma formation under oxygen-deficient conditions. Plant Cell 29, 775–790

Ye Y, Zhou L, Liu X, Liu H, Li D, Cao M, Chen H, Xu L, Zhu JK, Zhao Y. 2017. A novel chemical inhibitor of ABA signaling targets all ABA receptors. Plant Physiol 173, 2356–2369

Zhao Y, Chan Z, Xing L, Liu X, Hou YJ, Chinnusamy V, Wang P, Duan C, Zhu JK. 2013. The unique mode of action of a divergent member of the ABA-receptor protein family in ABA and stress signaling. Cell Res 23, 1380–1395

Zhou L, Duan J, Wang XM, Zhang HM, Duan MX, Liu JY. 2011. Characterization of a novel annexin gene from cotton (Gossypium hirsutum cv CRI 35) and antioxidative role of its recombinant protein. J Integr Plant Biol 53, 347–357

Zhu S, Yu X, Wang X, et al. 2007. Two calcium-dependent protein kinases, CPK4 and CPK11, regulate abscisic acid signal transduction in Arabidopsis. Plant Cell 19, 3019–3036

